# SARS-CoV-2 in Brazil: analysis of molecular variance and genetic diversity in viral haplotypes found in the states of Rio de Janeiro, São Paulo, Paraná and Tocantins

**DOI:** 10.1101/2020.12.02.409037

**Authors:** Rosane Maria de Albuquerque, Eduarda Doralice Alves Braz Da Silva, Dallynne Bárbara Ramos Venâncio, Robson da Silva Ramos, Pierre Teodósio Felix

**Author notes:** Corresponding author/ **Contact**.

## Abstract

In this work, 18 sequences of the SARS-CoV-2 virus were used, from four Brazilian states (Rio de Janeiro, São Paulo, Paraná and Tocantins) with 09, 04, 04, 8 and 01 haplotypes, respectively, with lengths ranging from 234 to 29,903 bp. All sequences were publicly available on the National Biotechnology Information Center (NCBI) platform and were previously aligned with the MEGA X software, where all gaps and ambiguous sites were extracted for the construction of the phylogenetic tree. Of the 301 sites analyzed, 68% varied, 131 of which were parsimonium-informative sites. Phylogenetic analyses revealed the presence of two distinct subgroups, corroborated by the high F_ST_ (80%). The high degree of polymorphism found among these samples helped to establish a clear pattern of non-genetic structuring, based on the time of divergence between the groups. All molecular variance estimators confirmed that there was no consensus in the conservation of the studied sequences, also indicating a high variation for the protein products of the virus. In a highly miscegenational and diverse population such as the Brazilian population, this observation draws our attention to the need for an urgent increase in public health actions, awareness strategies, hygiene and distancing practices and not the other way around.

## 1. Introduction

COVID-19 had its first case registered in Brazil in February 2020 and since then, the virus has expanded exponentially throughout Brazil. Part of this expansion is mainly due to the lack of infrastructure in the public health system (due to lack of personnel, medical equipment, diagnostic and/or therapeutic material) and misuse or even “non-use” of personal protective equipment (MILLER *et al*, 2020). With rapid dissemination and a diversified expansion in specific regions of Brazil, usually marked by insufficient sanitation and hygiene measures, SARS-CoV-2 has been establishing itself as the pandemic whose efforts in the use of protocols are by far the most inefficient since colonization (FELIX *et al*, 2020a).

Several factors such as: non-obedience to the use of masks, hand washing, the use of alcohol at 70%, an intense disagreement among members of the government about the practice of social isolation, a low population’s compliance with quarantine, the superstition about the virus infecting only people over 60 years of age, the lack of understanding of the population about the disease or the complete denial of its existence (inflated by denialist political leaders), from the early dismantling of the few existing field hospitals and finally the loss of the validity of thousands of RT-PCR tests stocked for a supposed distribution, only increase the rates of transmission, therefore the number of patients and unfortunately the number of deaths (FÉLIX *et al*, 2020b).

However, we at the Laboratory of Population Genetics and Computational Evolutionary Biology (LaBECom-UNIVISA) believe that contributing to genetic-population studies of SARS-CoV-2 in Brazil can be a strategy for the mitigation of infection, more precisely when trying to establish relationships between the diversity of the Brazilian population (highly miscigenous and genetically diverse) and the molecular diversity of the virus. This relationship would help us to assume, among other things, that the more conserved the viral sequences found in Brazil, the better the chances of resistance of the population, since the genetic diversity of the Brazilian people is also reflected, in a huge plasticity of clinical forms of COVID-19.

## 2. Objective

To evaluate the possible levels of polymorphism existing in 18 haplotypes of SARS-CoV-2 in Brazil.

## 3. Methodology

### Database

the 18 publicly available SARS-CoV-2 haplotypes from Brazil were rescued from the National Biotechnology Information Center (NCBI) platform at the address (https://www.ncbi.nlm.nih.gov/labs/virus/vssi/#/virus?SeqType_s=Nucleotide&VirusLineage_ss=Severe%20acute%20respiratory%20syndrome%20coronavirus%202,%20taxid:2697049&Country_s=Brazil) on November 27, 2020.

### Phylogenetic analyses

Nucleotide sequences previously described were used for phylogenetic analyses. The sequences were aligned using the MEGA X program (TAMURA *et al*., 2018) and the gaps were extracted for the construction of phylogenetic trees.

### Genetic Structuring Analyses

Paired F_ST_ estimators, Molecular Variance (AMOVA), Genetic Distance, mismatch, demographic and spatial expansion analyses, molecular diversity and evolutionary divergence time were obtained with the Software Arlequin v. 3.5 (EXCOFFIER *et al*., 2005) using 1000 random permutations (NEI and KUMAR, 2000). The F_ST_ and geographic distance matrices were not compared. All steps of this process are described below.

#### FOR GENETIC DIVERSITY

Among the routines of LaBECom, this test is used to measure the genetic diversity that is equivalent to the heterozygosity expected in the groups studied. We used for this the standard index of genetic diversity H, described by Nei (1987). Which can also be estimated by the method proposed by PONS and PETIT (1995).

#### FOR SITE FREQUENCY SPECTRUM (SFS)

According to LaBECom protocols, we used this local frequency spectrum analytical test (SFS), from DNA sequence data that allows us to estimate the demographic parameters of the frequency spectrum. Simulations are made using fastsimcoal2 software, available in http://cmpg.unibe.ch/software/fastsimcoal2/.

#### FOR MOLECULAR DIVERSITY INDICES

Molecular diversity indices are obtained by means of the average number of paired differences, as described by Tajima in 1993, in this test we used sequences that do not fit the model of neutral theory that establishes the existence of a balance between mutation and genetic drift.

#### FOR CALCULATING THETA ESTIMATORs

Theta population parameters are used in our Laboratory when we want to qualify the genetic diversity of the populations studied. These estimates, classified as Theta Hom – which aim to estimate the expected homozygosity in a population in equilibrium between drift and mutation and the estimates Theta (S) (WATTERSON, 1975), Theta (K) (EWENS, 1972) and Theta (π) (TAJIMA, 1983).

#### FOR THE CALCULATION OF The DISTRIBUTION OF MISMATCH

In LaBECom, analyses of the mismatch distribution are always performed relating the observed number of differences between haplotype pairs, trying to define or establish a pattern of population demographic behavior, as already described by (ROGERS; HARPENDING, 1992; Hudson, Hudson, HUDSON, SLATKIN, 1991; RAY et al., 2003, EXCOFFIER, 2004).

#### FOR PURE DEMOGRAPHIC EXPANSION

This model is always used when we intend to estimate the probability of differences observed between two haplotypes not recombined and randomly chosen, this methodology in our laboratory is used when we assume that the expansion, in a haploid population, reached a momentary balance even having passed through τ generations, of sizes 0 N to 1 N. In this case, the probability of observing the S differences between two non-recombined and randomly chosen haplotypes is given by the probability of observing two haplotypes with S differences in this population (Watterson, 1975).

#### FOR SPATIAL EXPANSION

The use of this model in LaBECom is usually indicated if the reach of a population is initially restricted to a very small area, and when one notices signs of a growth of the same, in the same space and over a relatively short time. The resulting population generally becomes subdivided in the sense that individuals tend to mate with geographically close individuals rather than random individuals. To follow the dimensions of spatial expansion, we at LaBECom always take into account:

L: Number of loci
Gamma Correction: This fix is always used when mutation rates do not seem uniform for all sites.
nd: Number of substitutions observed between two DNA sequences.
ns: Number of transitions observed between two DNA sequences.
nv: Number of transversions observed between two DNA sequences.
ω: G + C ratio, calculated in all DNA sequences of a given sample.
Paired Difference: Shows the number of loci for which two haplotypes are different.
Percentage difference: This difference is responsible for producing the percentage of loci for which two haplotypes are different.

#### FOR HAPLOTYPIC INFERENCES

We use these inferences for haplotypic or genotypic data with unknown gametic phase. Following our protocol, inferences are estimated by observing the relationship between haplotype i and xi times its number of copies, generating an estimated frequency (^pi). With genotypic data with unknown gametic phase, the frequencies of haplotypes are estimated by the maximum likelihood method, and can also be estimated using the expected Maximization (MS) algorithm.

#### FOR THE METHOD OF JUKES AND CANTOR

This method, when used in LaBECom, allows estimating a corrected percentage of how different two haplotypes are. This correction allows us to assume that there have been several substitutions per site, since the most recent ancestor of the two haplotypes studied. Here, we also assume a correction for identical replacement rates for all four nucleotides A C, G and T.

#### FOR KIMURA METHOD WITH TWO PARAMETERS

Much like the previous test, this fix allows for multiple site substitutions, but takes into account different replacement rates between transitions and transversions.

#### FOR TAMURA METHOD

We at LaBECom understand this method as an extension of the 2-parameter Kimura method, which also allows the estimation of frequencies for different haplotypes. However, transition-transversion relationships as well as general nucleotide frequencies are calculated from the original data.

#### FOR The TAJIMA AND NEI METHOD

At this stage, we were also able to produce a corrected percentage of nucleotides for which two haplotypes are different, but this correction is an extension of the Jukes and Cantor method, with the difference of being able to do this from the original data.

#### FOR TAMURA AND NEI MODEL

As in kimura’s models 2 parameters a distance of Tajima and Nei, this correction allows, inferring different rates of transversions and transitions, besides being able to distinguish transition rates between purines and pyrimidines.

#### FOR ESTIMATING DISTANCES BETWEEN HAPLOTYPES PRODUCED BY RFLP

We use this method in our laboratory when we need to verify the number of paired differences scouting the number of different alleles between two haplotypes generated by RFLP.

#### TO ESTIMATE DISTANCES BETWEEN HAPLOTYPES PRODUCED MICROSATELLITES

In this case, what applies is a simple count of the number of different alleles between two haplotypes. Using the sum of the square of the differences of repeated sites between two haplotypes (Slatkin, 1995).

#### MINIMUM SPANNING NETWORK

To calculate the distance between OTU (operational taxonomic units) from the paired distance matrix of haplotypes, we used a Minimum Spanning Network (MSN) tree, with a slight modification of the algorithm described in Rohlf (1973). We usually use free software written in Pascal called MINSPNET. EXE running in DOS language, previously available at: http://anthropologie.unige.ch/LGB/software/win/min-span-net/.

#### FOR GENOTYPIC DATA WITH UNKNOWN GAMETIC PHASE

##### EM algorithm

To estimate haplotypic frequencies we used the maximum likelihood model with an algorithm that maximizes the expected values. The use of this algorithm in LaBECom, allows to obtain the maximum likelihood estimates from multilocal data of gametic phase is unknown (phenotypic data). It is a slightly more complex procedure since it does not allow us to do a simple gene count, since individuals in a population can be heterozygous to more than one locus.

##### ELB algorithm

Very similar to the previous algorithm, ELB attempts to reconstruct the gametic phase (unknown) of multilocal genotypes by adjusting the sizes and locations of neighboring loci to explore some rare recombination.

#### FOR NEUTRALITY TESTS

##### Ewens-Watterson homozygosis test

We use this test in LaBECom for both haploid and diploid data. This test is used only as a way to summarize the distribution of allelic frequency, without taking into account its biological significance. This test is based on the sampling theory of neutral alleles from Ewens (1972) and tested by Watterson (1978). It is now limited to sample sizes of 2,000 genes or less and 1,000 different alleles (haplotypes) or less. It is still used to test the hypothesis of selective neutrality and population balance against natural selection or the presence of some advantageous alleles.

##### Accurate Ewens-Watterson-Slatkin Test

This test created by Slatikin in 1994 and adapted by himself in 1996. is used in our protocols when we want to compare the probabilities of random samples with those of observed samples.

##### Chakraborty’s test of population amalgamation

This test was proposed by Chakraborty in 1990, serves to calculate the observed probability of a randomly neutral sample with a number of alleles equal to or greater than that observed, it is based on the infinite allele model and sampling theory for neutral Alleles of Ewens (1972).

##### Tajima Selective Neutrality Test

We use this test in our Laboratory when DNA sequences or haplotypes produced by RFLP are short. It is based on the 1989 Tajima test, using the model of infinite sites without recombination. It commutes two estimators using the theta mutation as a parameter.

##### FS FU Test of Selective Neutrality

Also based on the model of infinite sites without recombination, the FU test is suitable for short DNA sequences or haplotypes produced by RFLP. However, in this case, it assesses the observed probability of a randomly neutral sample with a number of alleles equal to or less than the observed value. In this case the estimator used is θ.

#### FOR METHODS THAT MEASURE INTERPOPULATION DIVERSITY

##### Genetic structure of the population inferred by molecular variance analysis (AMOVA)

This stage is the most used in the LaBECom protocols because it allows to know the genetic structure of populations measuring their variances, this methodology, first defined by Cockerham in 1969 and 1973) and, later adapted by other researchers, is essentially similar to other approaches based on analyses of gene frequency variance, but takes into account the number of mutations between haplotypes. When the population group is defined, we can define a particular genetic structure that will be tested, that is, we can create a hierarchical analysis of variance by dividing the total variance into covariance components by being able to measure intra-individual differences, interindividual differences and/or interpopulation allocated differences.

##### Minimum Spanning Network (MSN) among haplotypes

In LaBECom, this tree is generated using the operational taxonomic units (OTU). This tree is calculated from the matrix of paired distances using a modification of the algorithm described in Rohlf (1973).

##### Locus-by-locus AMOVA

We performed this analysis for each locus separately as it is performed at the haplotypic level and the variance components and f statistics are estimated for each locus separately generating in a more global panorama.

##### Paired genetic distances between populations

This is the most present analysis in the work of LaBECom. These generate paired F_ST_ parameters that are always used, extremely reliably, to estimate the short-term genetic distances between the populations studied, in this model a slight algorithmic adaptation is applied to linearize the genetic distance with the time of population divergence (Reynolds et al. 1983; Slatkin, 1995).

##### Reynolds Distance (Reynolds et al. 1983)

Here we measured how much pairs of fixed N-size haplotypes diverged over t generations, based on F_ST_ indices.

##### Slatkin’s linearized F_ST’s_ (Slatkin 1995)

We used this test in LaBECom when we want to know how much two Haploid populations of N size diverged t generations behind a population of identical size and managed to remain isolated and without migration. This is a demographic model and applies very well to the phylogeography work of our Laboratory.

##### Nei’s average number of differences between populations

In this test we assumed that the relationship between the gross (D) and liquid (AD) number of Nei differences between populations is the increase in genetic distance between populations (Nei and Li, 1979).

##### Relative population sizes: divergence between populations of unequal sizes

We used this method in LaBECom when we want to estimate the time of divergence between populations of equal sizes (Gaggiotti and Excoffier, 2000), assuming that two populations diverged from an ancestral population of N0 size a few t generations in the past, and that they have remained isolated from each other ever since. In this method we assume that even though the sizes of the two child populations are different, the sum of them will always correspond to the size of the ancestral population. The procedure is based on the comparison of intra and inter populational (π’s) diversities that have a large variance, which means that for short divergence times, the average diversity found within the population may be higher than that observed among populations. These calculations should therefore be made if the assumptions of a pure fission model are met and if the divergence time is relatively old. The results of this simulation show that this procedure leads to better results than other methods that do not take into account unequal population sizes, especially when the relative sizes of the daughter populations are in fact unequal.

##### Accurate population differentiation tests

We at LaBECom understand that this test is an analog of fisher’s exact test in a 2×2 contingency table extended to a rxk contingency table. It has been described in Raymond and Rousset (1995) and tests the hypothesis of a random distribution of k different haplotypes or genotypes among r populations.

##### Assignment of individual genotypes to populations

Inspired by what had been described in Paetkau *et al* (1995, 1997) and Waser and Strobeck (1998) this method determines the origin of specific individuals, knowing a list of potential source populations and uses the allelic frequencies estimated in each sample from their original constitution.

##### Detection of loci under selection from F-statistics

We use this test when we suspect that natural selection affects genetic diversity among populations. This method was adapted by Cavalli-Sforza in 1996 from a 1973 work by Lewontin and Krakauer.

##### For tree construction using FIGTREE V 1.4.4. (VLAD *et al*, 2008)

To assemble molecular phylogeny, we used a number of 100 pseudo-replications for Bootstrap and as an evolutionary model, we used the 2-parameter Kimura model and as rates among sites a gamma distributed with invariant sites (G+I). We used nearest-neighbor-interchange (NNI) as ML Heuristic Method, the tree generated by MEGA X, was exported in *Newick* format and served as input for FIGTREE 1.4.4 software. In assembling the tree in FIGTREE, we assumed the need to display the subsequent probabilities of each of the clades present, as well as the estimation of the age of each node. We also designed a time scale axis for evolutionary history and for proper sizing of this time, we defined a reverse scale on the time axis. We chose to draw the tree with thick lines and color the clades by selecting the branches. Finally, we export the tree still in NEXUS and generate graphic files in PNG.

## 4. Results

### General properties of analyzed sequences

The 18 sequences studied came from four Brazilian states (São Paulo, Rio de Janeiro, Tocantins and Paraná) and had sizes ranging from 234 to 29,903 bp in length. Only a lot of 301 sites could be analyzed after alignment, revealing a percentage of 32% of conservation (78 sites). Among the 206 sites that varied, 64% (131 sites) revealed themselves as parsimonium-informative, evidencing the high degree of polymorphism for the whole studied set. The graphical representation of these sites could be seen in a logo built with the WEBLOGO 3 program (CROOKS *et al*., 2004), where the size of each nucleotide is proportional to its frequency for certain sites. (Figure 1).

**Figure 1:**
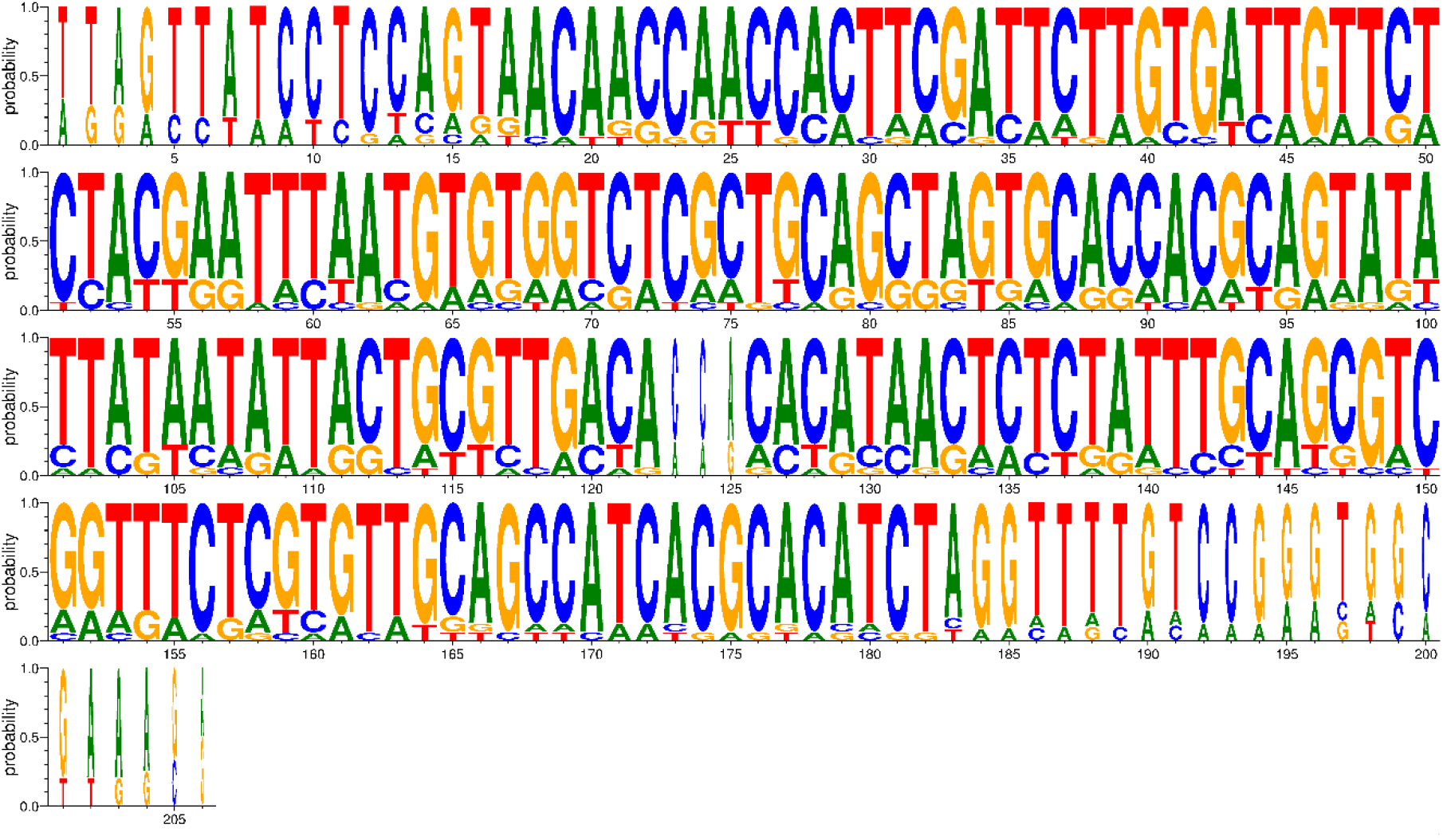
Graphic representation of the 206 variable sites for the 18 haplotypes of SARS-CoV-2 in Brazil. * Generated by WEBLOGO v.3.0 (CROOKS *et al*., 2004)

Using the Maximum Likelihood method, based on the 131 parsimony-informative sites, it was possible to understand that the 18 haplotypes comprised two distinct subgroups, one of which is composed exclusively of the samples from the State of Tocantins. The phylogenetic relationships for the whole set, as well as the divergence time found between all sequences, suggest a high degree of differentiation for the virus between the North and Southeast regions of Brazil (Figure 2 and Figure 3).

**Figure 2.**
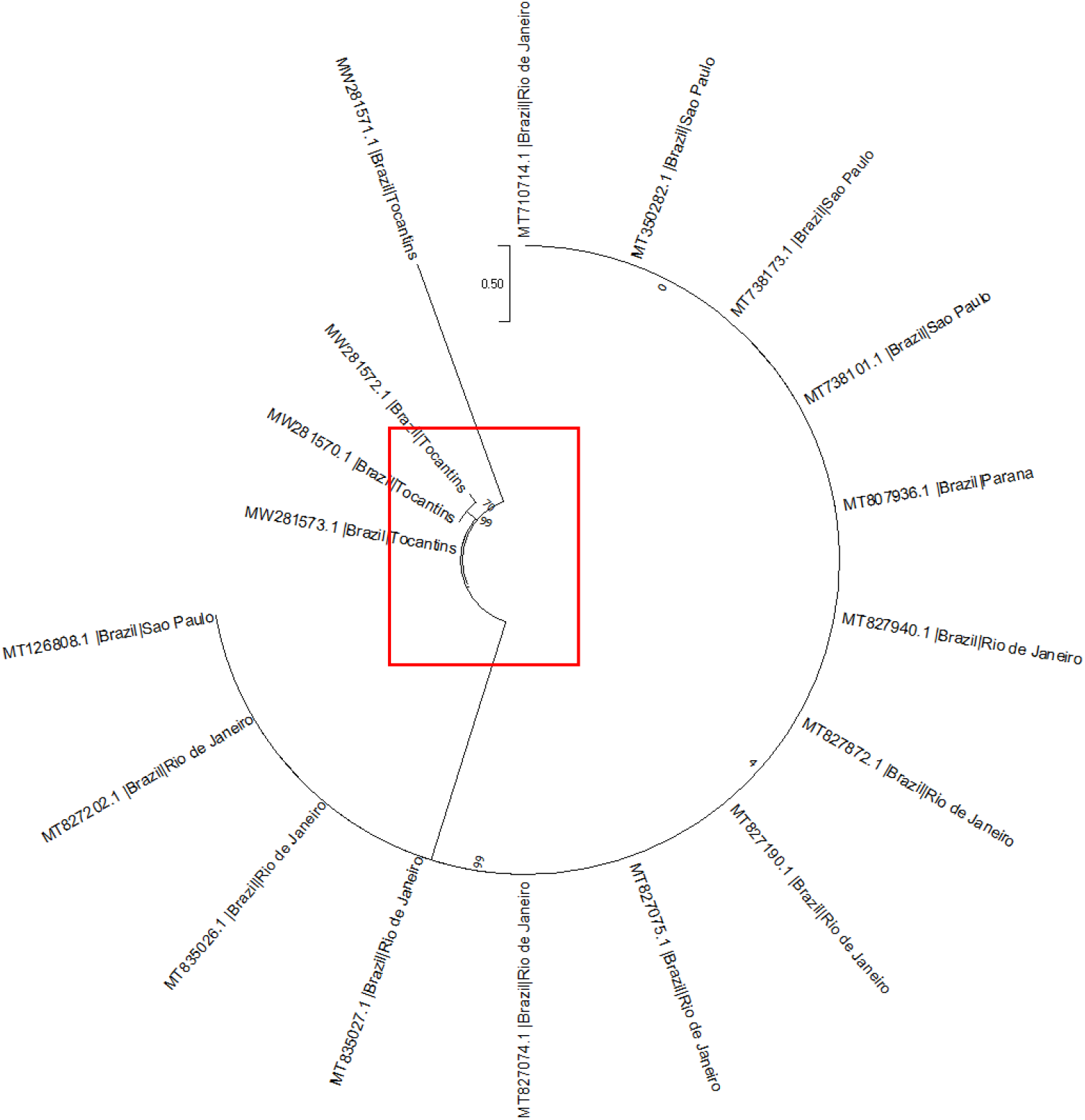
Evolutionary analysis by Maximum Likelihood method. The evolutionary history was inferred by using the Maximum Likelihood method and Tamura 3-parameter model [1]. The tree with the highest log likelihood (−1106.62) is shown. The percentage of trees in which the associated taxa clustered together is shown next to the branches. Initial tree(s) for the heuristic search were obtained automatically by applying Neighbor-Join and BioNJ algorithms to a matrix of pairwise distances estimated using the Tamura 3 parameter model, and then selecting the topology with superior log likelihood value. A discrete Gamma distribution was used to model evolutionary rate differences among sites (5 categories (+*G*, parameter = 62.9618)). The rate variation model allowed for some sites to be evolutionarily invariable ([+*I*], 8.43% sites). This analysis involved 18 nucleotide sequences. Codon positions included were 1st+2nd+3rd+Noncoding. There were a total of 301 positions in the final dataset. Evolutionary analyses were conducted in MEGA X [2]

**Figure 3.**
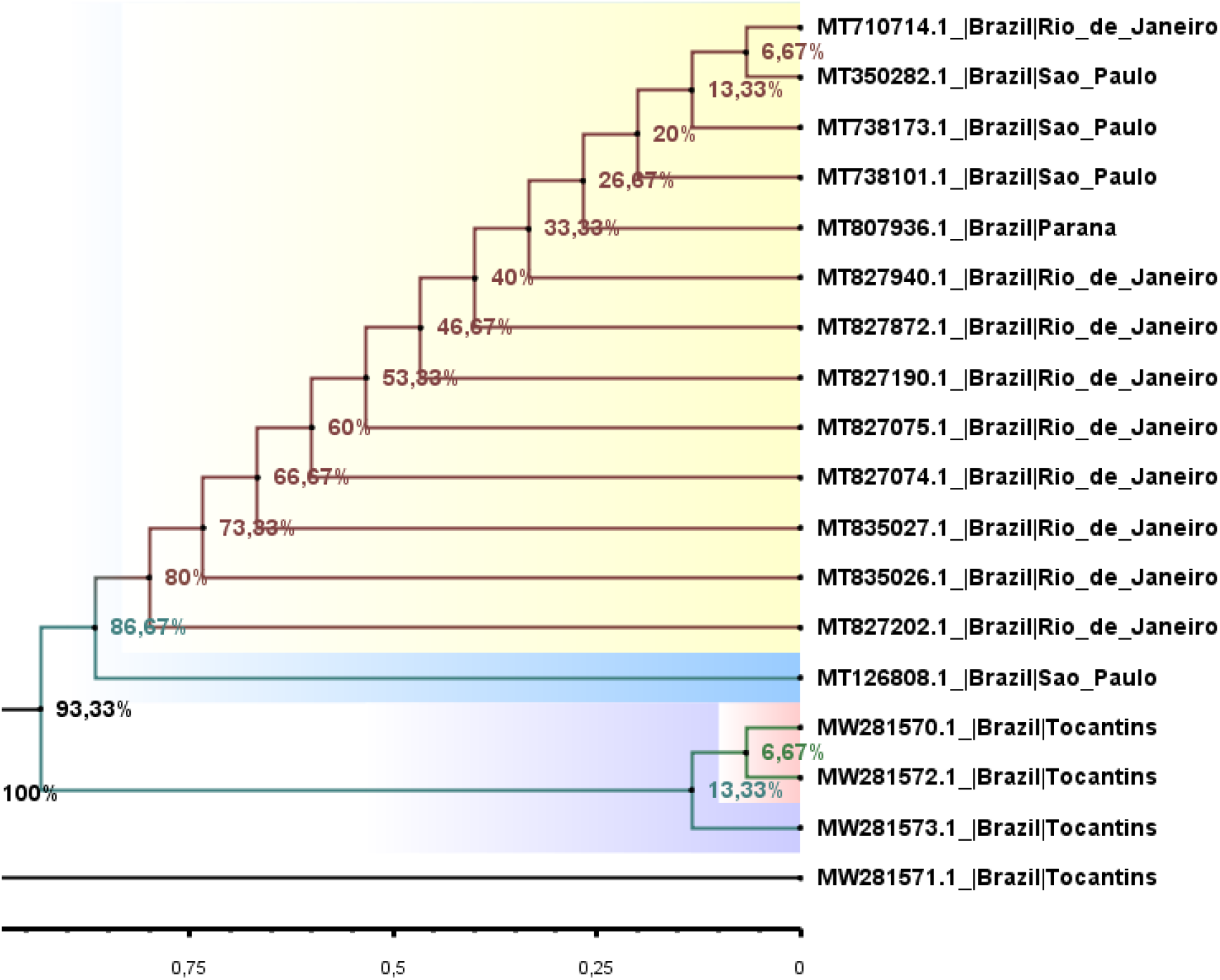
Figure 3. Phylogenetic Tree of 18 Sequences of SARS-CoV-2 from 04 Brazilian states. The main geographic groupings of SARS-CoV-2 are indicated and the subsequent probability values are shown for key nodes. In all cases, the times of evolutionary divergence correspond to the percentages shown. **Note:** small clusters formed for haplotypes of different localities (colors). * Figure generated by FigTree software version V 1.4.4.

### Genetic Distance Analysis

The analyses based on F_ST_ values also confirmed the presence of two distinct genetic “entities” with components of variation greater than 80% and with significant evolutionary divergences for Tocantins samples, as well as a high genetic similarity between the sequences that comprised the Southeast and South regions of Brazil, for a *p-*value lower than 0.05 (table 1).

**Table 1.** Components of haplotypic variation and paired F_ST_ value for the 18 haplotypes of SARS-CoV-2 in Brazil

| Source of variation | d.f. | Sum of squares | Variance components | Percentage of variation |
| --- | --- | --- | --- | --- |
| Among populations | 3 | 258.611 | 20.87177 Va | 80.56 |
| Within populations | 14 | 70.500 | 5.03571 Vb | 19.44 |
| Total | 17 | 329.111 | 25.90748 |  |
| Fixation Index | **$F_{ST}$:** | **0.80563** |  |  |
| Significance tests (1023 permutations) |  |  |  |  |
| Va and $F_{ST}$: P (rand. value > obs. value) = 0.00000 | | | | |
| P (rand. value = obs. value) = 0.00000 |  |  |  |  |
| P-value = 0.00000+-0.00000 |  |  |  |  |

The analyses also confirmed a high genetic dissimilarity among all haplotypes. However, the use of the divergence matrix in the construction of the tree helped in the recognition of minimal similarities between some haplotypes, including at different geographical points. The maximum divergence patterns were also obtained when less robust methods of phylogenetic pairing (e.g. UPGMA) were used, reflecting the non-haplotypic structure in the clades. With the use of a divergence matrix, it was possible to identify geographical variants that had shorter genetic distances and the “*a posteriori*” probabilities were able to separate the main clusters into additional small groups, confirming the presence of a minimum probability of kinship between haplotypes. The *Tau* variations (99%), related to the ancestry of the two subgroups, revealed a significant time of divergence, supported by mismatch analysis and demographic and spatial expansion analyses (Figure 4, Figure 5, Table 2).

**Figure 4.**
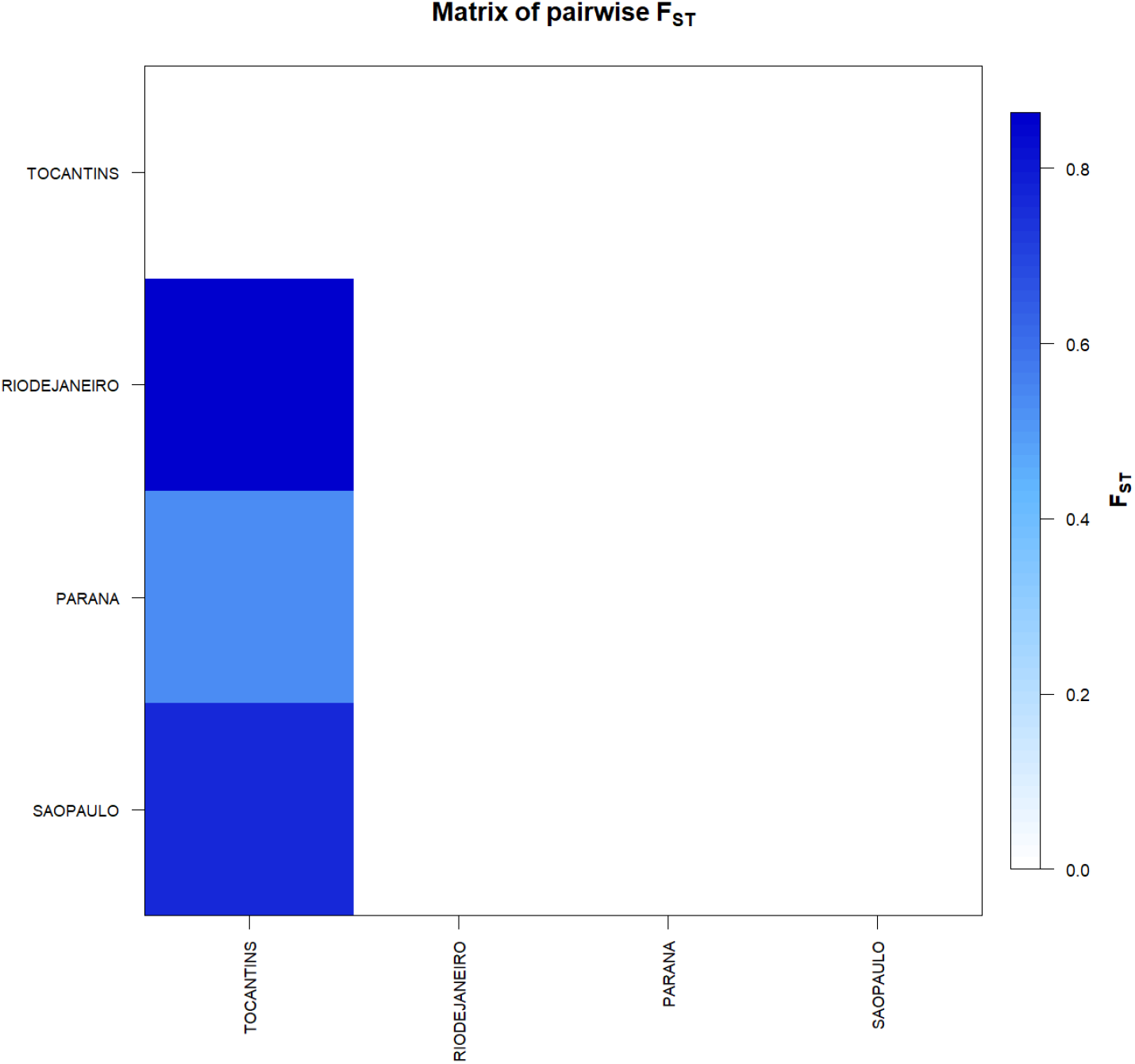
F_ST_-based genetic distance matrix between for the 18 haplotypes of SARS-CoV-2 in Brazil. * Generated by the statistical package in R language using the output data of the Software Arlequin version 3.5.1.2

**Figure 5.**
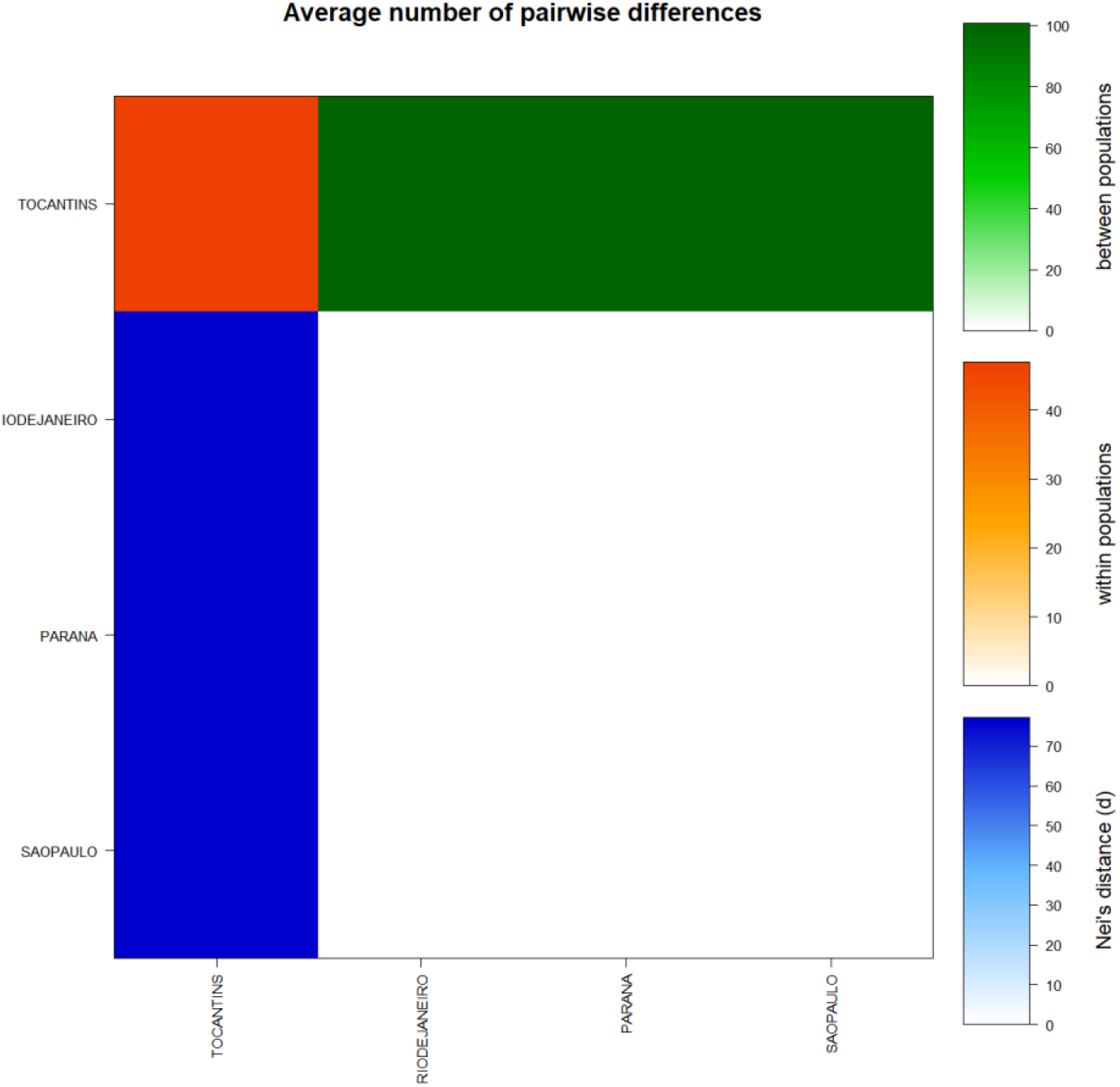
Matrix of paired differences between the populations studied: between the groups; within the groups; and Nei distance for the 18 haplotypes of SARS-CoV-2 in Brazil.

**Table 2.** Demographic and spatial expansion simulations based on the τ, θ, and M indices of sequences of the 18 haplotypes of SARS-CoV-2 in Brazil.

| Statistics | TOCANTINS | RIO DE JANEIRO | PARANÁ | SÃO PAULO | Mean | s.d. |
| --- | --- | --- | --- | --- | --- | --- |
| **Demographic expansion** |  |  |  |  |  |  |
| ----- |  |  |  |  |  |  |
| Tau | 0.00000 | 0.00000 | 0.00000 | 0.00000 | 0.00000 | 0.00000 |
| Theta0 | 0.40018 | 0.00000 | 0.00000 | 0.00000 | 0.10004 | 0.20009 |
| Theta1 | 6827.33131 | 0.00000 | 0.00000 | 0.00000 | 1706.83283 | 3413.66566 |
| SSD | 0.34125 | 0.00000 | 0.00000 | 0.00000 | 0.08531 | 0.17063 |
| Raggedness index | 0.75000 | 0.00000 | 0.00000 | 0.00000 | 0.18750 | 0.37500 |
| **Spatial expansion** |  |  |  |  |  |  |
| ----- |  |  |  |  |  |  |
| Tau | 99.10791 | 0.00000 | 0.00000 | 0.00000 | 24.77698 | 49.55396 |
| Theta | 0.00072 | 0.00000 | 0.00000 | 0.00000 | 0.00018 | 0.00036 |
| M | 1.05501 | 0.00000 | 0.00000 | 0.00000 | 0.26375 | 0.52751 |
| SSD | 0.23307 | 0.00000 | 0.00000 | 0.00000 | 0.05827 | 0.11653 |
| Raggedness index | 0.75000 | 0.00000 | 0.00000 | 0.00000 | 0.18750 | 0.37500 |

### Molecular diversity analyses

Molecular diversity analyses estimated by θ, in the Tocantins, reflected a significant level of mutations among all haplotypes (transitions and transversions). Indel mutations (insertions or deletions) were not found in either of the two groups studied. The D tests of Tajima and Fs de Fu showed disagreements between the estimates of general θ and π, but with negative and highly significant values, indicating an absence of population expansion. The irregularity index (R= Raggedness) with parametric bootstrap simulated new values θ for before and after a supposed demographic expansion and in this case assumed a value equal to zero for the groups (table 3 and table 4 and figure 6).

**Table 3.** Molecular Diversity Indices for the 18 haplotypes of SARS-CoV-2 in Brazil

| Statistics | TOCANTINS | RIO DE JANEIRO | PARANÁ | SÃO PAULO | Mean | s.d. |  |
| --- | --- | --- | --- | --- | --- | --- | --- |
| No. of transitions | 44 | 0 | 0 | 0 | 11.000 | 22.000 |  |
| No. of transversions | 53 | 0 | 0 | 0 | 13.250 | 26.500 |  |
| No. of substitutions | 97 | 0 | 0 | 0 | 24.250 | 48.500 |  |
| No. of indels | 0 | 0 | 0 | 0 | 0.000 | 0.000 |  |
| No. of ts. sites | 44 | 0 | 0 | 0 | 11.000 | 22.000 |  |
| No. of tv. sites | 53 | 0 | 0 | 0 | 13.250 | 26.500 |  |
| No. of subst. sites | 97 | 0 | 0 | 0 | 24.250 | 48.500 | Total:97 |
| No. private subst. sites | 108 | 0 | 0 | 0 | 27.000 | 54.000 |  |
| No. of indel sites | 0 | 0 | 0 | 0 | 0.000 | 0.000 |  |
| Pi | 48.500 | 0.000 | 0.000 | 0.000 | 12.12500 | 24.25000 |  |
| Theta_k | N.A. | N.A. | N.A. | N.A. | N.A. | N.A. |  |
| Theta_k_lower | N.A. | N.A. | N.A. | N.A. | N.A. | N.A. |  |
| Theta_k_upper | N.A. | N.A. | N.A. | N.A. | N.A. | N.A. |  |
| Theta_H | N.A. | N.A. | N.A. | N.A. | N.A. | N.A. |  |
| s.d. Theta_H | N.A. | N.A. | N.A. | N.A. | N.A. | N.A. |  |
| Theta_S | 52.90909 | 0.00000 | 0.00000 | 0.00000 | 13.22727 | 26.45455 |  |
| s.d. Theta_S | 28.76489 | 0.00000 | 0.00000 | 0.00000 | 7.19122 | 14.38245 |  |
| Theta_pi | 48.50000 | 0.00000 | 0.00000 | 0.00000 | 12.12500 | 24.25000 |  |
| s.d. Theta_pi | 32.07536 | 0.00000 | 0.00000 | 0.00000 | 8.01884 | 16.03768 |  |

**Table 4.** Neutrality Tests for the 18 haplotypes of SARS-CoV-2 in Brazil

|  | Statistics | TOCANTINS | RIO DE JANEIRO | PARANA | SÃO PAULO | Mean | s.d. |
| --- | --- | --- | --- | --- | --- | --- | --- |
| **Ewens-Watterson test** |  |  |  |  |  |  |  |
|  | Sample size | 4 | 9 | 1 | 4 | 4.50000 | 3.31662 |
|  | No. of alleles(unchecked) | 4 | 9 | 1 | 4 | 4.50000 | 3.31662 |
|  | Observed F value | N.A. | N.A. | N.A. | N.A. | N.A. | N.A. |
|  | Expected F value | N.A. | N.A. | N.A. | N.A. | N.A. | N.A. |
|  | Watterson test: Pr(rand F <= obs F) | N.A. | N.A. | N.A. | N.A. | N.A. | N.A. |
|  | Slatkin's exact test P-value | N.A. | N.A. | N.A. | N.A. | N.A. | N.A. |
| **Chakraborty's test** |  |  |  |  |  |  |  |
|  | Sample size | 4 | 9 | 1 | 4 | 4.50000 | 3.31662 |
|  | No. of alleles(unchecked) | 4 | 9 | 1 | 4 | 4.50000 | 3.31662 |
|  | Obs. homozygosity | 0.00000 | 0.00000 | 0.00000 | 0.00000 | 0.00000 | 0.00000 |
|  | Exp. no. of alleles | 3.88194 | 1.00000 | 0.00000 | 1.00000 | 1.47049 | 1.67533 |
|  | P(k or more alleles) | N.A. | N.A. | N.A. | N.A. | N.A. | N.A. |
| **Tajima's D test** |  |  |  |  |  |  |  |
|  | Sample size | 4 | 9 | 1 | 4 | 4.50000 | 3.31662 |
|  | S | 97 | 0 | 0 | 0 | 24.25000 | 48.50000 |
|  | Pi | 48.50000 | 0.00000 | 0.00000 | 0.00000 | 12.12500 | 24.25000 |
|  | Tajima's D | -0.87170 | 0.00000 | 0.00000 | 0.00000 | -0.21793 | 0.43585 |
|  | Tajima's D p-value | 0.01200 | 1.00000 | 1.00000 | 1.00000 | 0.75300 | 0.49400 |
| **Fu's FS test** |  |  |  |  |  |  |  |
|  | No. of alleles(unchecked) | 4 | 9 | 1 | 4 | 4.50000 | 3.31662 |
|  | Theta_pi | 48.50000 | 0.00000 | 0.00000 | 0.00000 | 12.12500 | 24.25000 |
|  | Exp. no. of alleles | 3.88194 | 1.00000 | 0.00000 | 1.00000 | 1.47049 | 1.67533 |
|  | FS | 2.05229 | 0.00000 | 0.00000 | 0.00000 | 0.00000 | 0.00000 |
|  | FS p-value | 0.50700 | 1.00000 | N.A. | 1.00000 | N.A. | N.A. |

**Figure 6.**
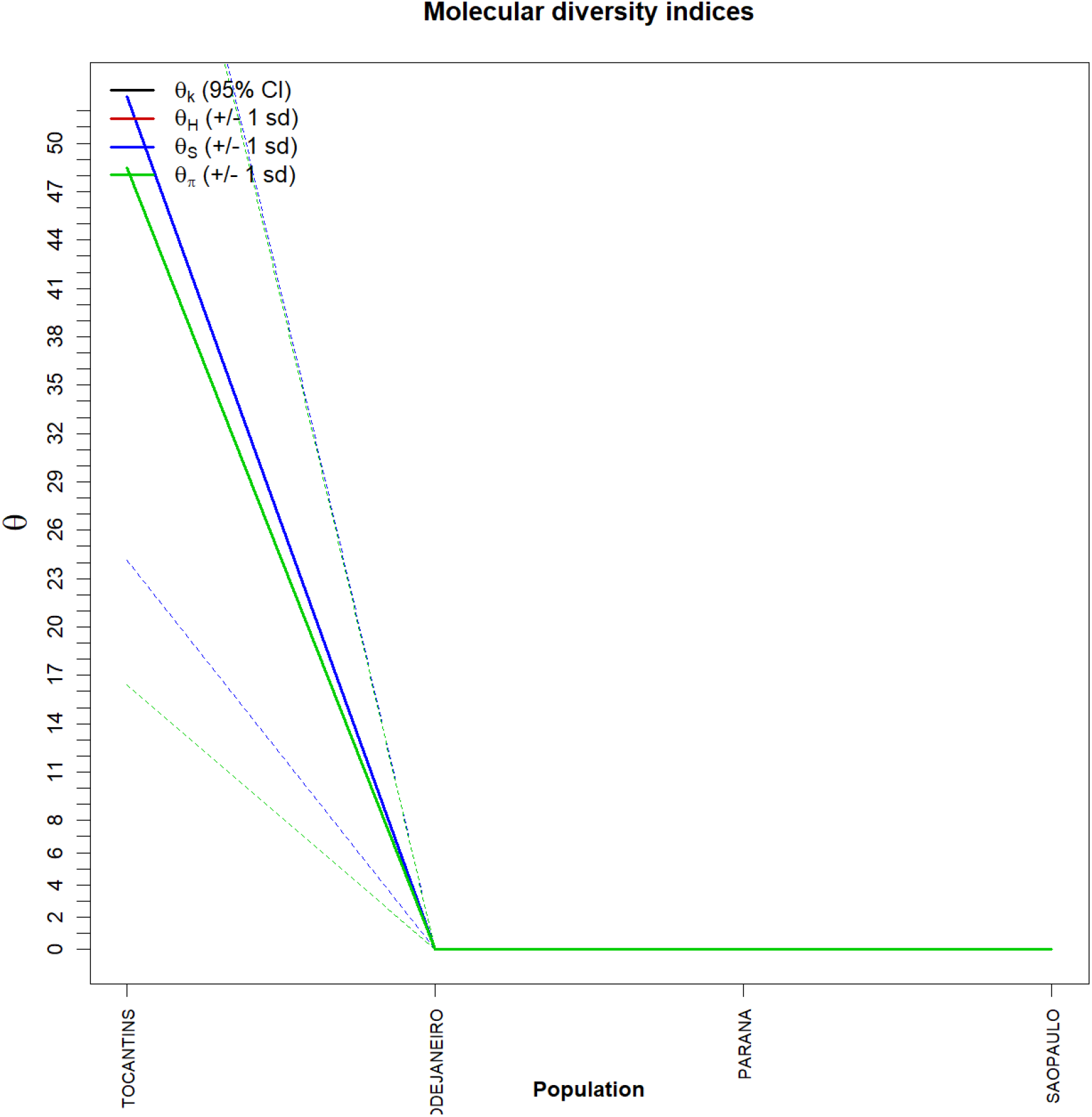
Graph of molecular diversity indices for the 18 haplotypes of SARS-CoV-2 in Brazil. In the graph the values of θ: (θk) Relationship between the expected number of alllos (k) and the sample size; (θH) Expected homozygosity in a balanced relationship between drift and mutation; (θS) Relationship between the number of segregating sites (S), sample size (n) and non-recombinant sites; (θπ) Relationship between the average number of paired differences (π) and θ. * Generated by the statistical package in R language using the output data of the Arlequin software version 3.5.1.2.

## 5. Discussion

With the use of phylogenetic analysis methodologies and population structure, it was possible to detect the existence of a small degree of similarity between the haplotypes of SARS-CoV-2 in Brazil. Because significant levels of structuring were not found, we assumed that there were high levels of variation probably related to a gain of intermediate haplotypes over time, associated, perhaps, with a significant increase in gene flow, especially in the states that make up the Southeast and South regions of Brazil. The occurrence of geographical isolations from the absence of past events of defragmentation may have generated this discontinuous pattern of genetic divergence between the groups, since the high values found for genetic distance support the presence of this pattern of divergence between haplotypes, as well as in the high frequency of polymorphisms. This suggests that molecular diversity may be due to non-synonymous substitutions as the main components of variations. All analyses supported that the data are a confirmation that there is no consensus in the conservation of the SARS-CoV-2 genome in Brazil, corroborating what was described by FELIX *et al.*, 2020a, for the Countries of South America, and it is therefore safe to affirm that the genetic variability of the Virus is different in different subsets for other Brazilian regions.

These considerations were also supported by simple phylogenetic pairing methodologies, such as UPGMA, which in this case, with a discontinuous pattern of genetic divergence between the groups, observed a large number of branches with many mutational stages. These mutations have possibly established themselves to drift due to the founding effect, which accompanies the dispersive behavior and/or loss of intermediate haplotypes throughout the generations. The values found for the genetic distance considered the minimum differences between the groups, as well as the inference of values greater than or equal to those observed in the proportion of these permutations, including the p-value of the test.

The discrimination of the two genetic entities (two subgroups) in distinct regions of The Brazilian territory was also perceived when the interhaplotypic variations were hierarchised in all components of covariance: by their intra and inter-individual differences or by their intra and intergroup differences, generating dendrograms that supported the idea that the significant differences found in the Tocantins group, for example, can even be shared in their form, but not in their number, since the result of estimates of the average evolutionary divergence within the Southeast and South groups were so low.

Since no relationship between genetic distance and geographic distance (mantel test) was made in this study, we assumed that the absence of genetic flow (observed by non-haplotypic sharing) between the studied regions, is due to the presence of natural geographical barriers. The φ estimators, although extremely sensitive to any form of molecular variation (Fu, 1997), supported the uniformity between the results found by all the methodologies employed, and can be interpreted as a phylogenetic confirmation that there is no consensus in the conservation of the studied sequences, and it is safe to even affirm that the large number of enontradepolymorphs should be reflited directly in a high variation for protein products. These considerations ensure that some responses to the different clinical forms of COVID-19 are not as efficient for the Brazilian population (highly misicigenated), a fact that draws attention to the urgent need for an increase and not the other way around, of public health actions, in awareness strategies, in hygiene practices and in social distancing.

## References

Cavalli-Sforza LL. Population structure and human evolution. Proc R Soc Lond B Biol Sci 164, 362–379. 1966.

Chakraborty, R. Mitochondrial DNA polymorphism reveals hidden heterogeneity within some Asian populations. Am. J. Hum. Genet. 47:87–94. 1990.

Cockerham, C. C., Analysis of gene frequencies. Genetics 74: 679–700. 1973.

Cockerham, C. C., Variance of gene frequencies. Evolution 23: 72–83. 1969.

Crooks G.E., Hon G., Chandonia J.M. Brenner S.E. WebLogo: A sequence logo generator, Genome Research, 14:1188–1190, 2004.

Ewens, W.J. The sampling theory of selectively neutral alleles. Theor. Popul. Biol. 3:87–112. 1972.

Excoffier L. Patterns of DNA sequence diversity and genetic structure after a range expansion: lessons from the infinite-island model. 2004.

Felix, P.T; Nascimento Filho, C.B.; Ramos, R.S.; Paulino, A.J.; Venâncio, D.B.R. Levels of genetic diversity of SARS-CoV-2 virus: reducing speculations about the genetic variability of the virus in South America. bioRxiv 2020a.

Felix, P.T; Nascimento Filho, C.B.; Ramos, R.S.; Paulino, A.J.; Venâncio, D.B.R. Molecular variance analysis (AMOVA) and levels of genetic diversity of complete genome of SARS-CoV-2 virus from of six South American Countries. Research square.2020b.

Fu, Y.X. Statistical tests of neutrality of mutations against population growth, hitchhiking and background selection. Genetics 147: 915–925. 1997.

Gaggiotti, O., and L. Excoffier. A simple method of removing the effect of a bottleneck and unequal population sizes on pairwise genetic distances. Proceedings of the Royal Society London B 267: 81–87. 2000.

Hudson, R. R. Gene genealogies and the coalescent process, pp. 1–44 in Oxford Surveys in Evolutionary Biology, edited by Futuyama, and J. D. Antonovics. Oxford University Press, New York. 1990.

Kumar S, Stecher G, Li M, Knyaz C; Tamura K. MEGA X: Molecular Evolutionary Genetics Analysis across computing platforms. Molecular Biology and Evolution 35:1547–1549. 2018.

Miller MJ, Loaiza JR, Takyar A, Gilman RH. COVID-19 na América Latina: Nova dinâmica de transmissão para uma pandemia global. PLoS Negl Trop Dis. 2020.

Nei M. and Kumar S. Molecular Evolution and Phylogenetics. Oxford University Press, New York.2000.

Nei, M. and W. H. Li. Mathematical model for studying genetic variation in terms of restriction endonucleases. Proc.Natl.Acad.Sci.USA 76:5269–5273.1979.

Nei, M. Molecular Evolutionary Genetics. Columbia University Press, New York, NY, USA.1987.

Paetkau D, Calvert W, Stirling I and Strobeck C. Microsatellite analysis of population structure in Canadian polar bears. Mol Ecol 4:347–54.1995.

Paetkau D, Waits LP, Clarkson PL, Craighead L and Strobeck C. An empirical evaluation of genetic distance statistics using microsatellite data from bear (Ursidae) populations. Genetics 147:1943–1957.1997.

Pons, O.; Petit, J.R. Estimation, Variance and Optimal Sampling of Gene Diversity I. Haploid locus. Theor Appl Genet 90: 462–470, 1995.

Ray N, Currat M, Excoffier L. Intra-Deme Molecular Diversity in Spatially Expanding Populations. Mol Biol Evol 20(1): 76–86.2003.

Raymond M. and F. Rousset. An exact tes for population differentiation. Evolution 49:1280–1283. 1995.

Reynolds, J., Weir, B.S., and Cockerham, C.C. Estimation for the coancestry coefficient: basis for a short-term genetic distance. Genetics 105:767–779. 1983.

Rogers, A. R., and H. Harpending, Population growth makes waves in the distribution of pairwise genetic differences. Mol. Biol. Evol. 9: 552–569.1992.

Rohlf, F. J., Algorithm 76. Hierarchical clustering using the minimum spanning tree. The Computer Journal 16:93–95.1973.

Slatkin, M. A measure of population subdivision based on microsatellite allele frequencies. Genetics 139: 457–462.1991.

Slatkin, M. A measure of population subdivision based on microsatellite allele frequencies. Genetics 139: 457–462.1995.

Tajima, F. Evolutionary relationship of DNA sequences in finite populations. Genetics 105: 437–460. 1983.

Tamura K. Estimation of the number of nucleotide substitutions when there are strong transition-transversion and G + C-content biases. Molecular Biology and Evolution 9:678–687. (1992).

Vlad I. Morariu, Balaji Vasan Srinivasan, Vikas C. Raykar, Ramani Duraiswami, and Larry S. Davis. Automatic online tuning for fast Gaussian summation. Advances in Neural Information Processing Systems (NIPS), 2008.

Waser PM, and Strobeck C. Genetic signatures of interpopulation dispersal. TREE.43–44.1998.

Watterson, G. The homozygosity test of neutrality. Genetics 88:405–417.1978.

Watterson, G., On the number of segregating sites in genetical models without recombination. Theor.Popul.Biol.7: 256–276.1975.

